# Complementary roles of orbitofrontal and prelimbic cortices in adaptation of reward motivated actions to learned anxiety

**DOI:** 10.1101/2023.08.17.553761

**Authors:** David S. Jacobs, Alina P. Bogachuk, Bita Moghaddam

## Abstract

**Background:** Anxiety is a common symptom of several mental health disorders and adversely affects motivated behaviors. Anxiety can emerge from associating risk of future harm while engaged in goal-guided actions. Using a recently developed behavioral paradigm to model this aspect of anxiety, we investigated the role of two cortical subregions, the prelimbic medial frontal cortex (PL) and lateral orbitofrontal cortex (lOFC), which have been implicated in anxiety and outcome expectation, in flexible representation of actions associated with harm risk.

**Methods:** A seek-take reward-guided instrumental task design was used to train animals to associate the seek action with a variable risk of punishment. After learning, animals underwent extinction training for this association. Fiber photometry was used to measure and compare neuronal activity in PL and lOFC during learning and extinction.

**Results:** Animals increased action suppression in response to punishment contingencies. This increase dissipated after extinction training. These behavioral changes were associated with region specific changes in neuronal activity. PL neuronal activity preferentially adapted to threat of punishment whereas lOFC activity adapted to safe aspects of the task. Moreover, correlated activity between these regions was suppressed during actions associated with harm risk suggesting that these regions may guide behavior independently under anxiety.

**Conclusions:** These findings suggest the PL and lOFC serve distinct but complementary roles in the representation of learned anxiety. This dissociation may provide a mechanism for how overlapping cortical systems are implicated in reward-guided action execution during anxiety.

## Introduction

Mammalian frontal and prefrontal cortex (PFC) subregions play a crucial role in action selection and execution that is driven by external and internal states. The influence of the PFC is especially critical when the appropriate or optimal action is novel or uncertain, and the inappropriate response is habitual or impulsive (Soltani & Koechlin, 2022; Yoshida & Ishii, 2006). Several PFC subregions have been extensively implicated in psychiatric illnesses including affective and addictive disorders (Milad & Rauch, 2007; Torregrossa et al., 2008; Volkow et al., 2003). A common symptom of these disorders is anxiety, which has been intimately associated with aberrant PFC activity (Balderston, Vytal, et al., 2017; Bishop, 2009; Brady et al., 2013; Roberts, 2020). While broad outlines of PFC subregion contributions to behavior have emerged, little is known about distinct functions of each of these cortical domains in the context of anxiety during action execution.

Anxiety is an affective state that supports behavioral adaptation to the presence of threat. While anxiety is a conserved and potentially evolutionarily useful process, it can lead to impaired decision making, maladaptive avoidance, and anxiety disorders (Kenwood, Kalin, et al., 2022). Anxiety is also a key cause of relapse in drug and alcohol use disorders and a debilitating symptom of PTSD and other mood disorders (Sinha, 2007). Animal models of anxiety have traditionally focused on innate behaviors and exploration of a novel context (Lezak et al., 2017). These approaches are operationally distinct from real world anxiety which often involves dread about a distant threat related to goal-guided behavior. These distant events are often associated with low or variable harm probability, with the threat association generally being learned and not innate. For example, anxiety about car or airplane crashes are learned, and can cause anxiety despite the low probability of the occurrence of these events. We have recently designed and characterized rodent behavioral paradigms modeling this form of anxiety where animals learn that reward-guided actions are associated with a risk of harm (Jacobs & Moghaddam, 2020; Park & Moghaddam, 2017b). The aim of the present study was to use this model of learned anxiety to understand the neuronal dynamics of two PFC subregions as animals learn that actions are associated with varying risk of harm. These regions included lateral orbitofrontal cortex (lOFC) and prelimbic medial PFC (PL). These regions are implicated in flexible planning of action selection and threat discrimination (Capuzzo & Floresco, 2020; Del Arco et al., 2017; Hardung et al., 2017; Ray et al., 2018; Sarlitto et al., 2018), and exhibit different roles in avoidance behavior, threat sensitivity (Diehl et al., 2018; Jean-Richard-dit-Bressel & McNally, 2016; Rodriguez-Romaguera et al., 2016), and rule encoding under pharmacological anxiety (Park et al., 2016).

We recorded neuronal calcium activity by using fiber photometry in lOFC and PL of male and female rats as they learned that a reward-guided action is associated with varying risk of punishment and subsequently learned that all actions were safe (extinction). We found distinct changes in neuronal activity between the PL and lOFC over the learning of risk of punishment. Neuronal activity changed in PL when actions were associated with threat of punishment whereas lOFC activity increased in response to actions which were safe. Furthermore, correlated activity between these regions were dynamically suppressed during actions associated with threat suggesting that these regions may guide action processing independently under anxiety.

## Materials & Methods

See supplemental methods for detailed surgical, behavioral, and fiber photometry procedures.

### Subjects

A total of 8 adult Long-Evans rats, housed on a reverse 12 h:12 h light/dark cycle, were used. Both males (n=6) and females (n=2) were utilized. All experimental procedures were approved by the Oregon Health and Science University Institutional Animal Care and Use Committee.

### Surgery

#### Viral Infusion and Optical Fiber Implant Surgery

To allow for pan-neuronal expression of fluorescent calcium indicator GCaMP6s as well as a non-calcium dependent fluorophore tdTomato, rats were injected with a viral construct (AAV8-hSyn-GCaMP6s-p2a-tdTomato) under anesthesia in the PL and lOFC. In a second surgery, at least 4 weeks later, an optical fiber was placed in the PL and lOFC and secured to the skull to permit recording of GCaMP activity.

### Behavioral Testing

#### Phase I – Reward seeking under safe conditions

Animals were trained to perform successive nosepokes to receive a sucrose pellet (45 mg) in an operant chamber according to previously published methods (Jacobs & Moghaddam, 2020). Briefly, completion of the first ‘seek’ action led to a 1.5 sec delay followed by illumination of the opposite ‘take’ nosepoke. A nosepoke on the illuminated ‘take’ nosepoke would then lead to a 1 sec delay followed by delivery of a sucrose pellet. Sessions were split into four 12 min blocks separated by a 2 min inter-block interval. No risk of shock was present for any action.

#### Phase II - Reward seeking under learning of threat of punishment

We utilized the Probabilistic Punishment Task in accordance with previously published methods (Jacobs & Moghaddam, 2020). Like safe sessions in Phase I, Phase II sessions consisted of four blocks of 20 trials each. However, there was a variable risk of receiving a punishment (300 ms electrical footshock at 0.25 mA) after the seek action which increased over the blocks (0%, 6%, 10%, and 18% in blocks 1, 2, 3, and 4, respectively). The take action-reward contingency remained safe and always produced reward.

#### Phase III - Extinction of threat of punishment during reward seeking

Following learning of the Probabilistic Punishment Task in Phase II, subjects underwent two extinction sessions where the risk of punishment was removed so all actions were safe.

#### Unsignaled footshock test

To determine if any learning related changes in footshock responsivity were products of footshock exposure, we applied the same footshock in a different context where subjects received six unsignaled footshocks after Phase III was completed.

### Fiber Photometry Recording

Recordings were performed with a commercially available fiber photometry system (Tucker-Davis Technologies RZ5). Recording was accomplished by providing both 470 nm and 560 nm excitation light to the PL and lOFC. Fluorescence was collected through dichroic mirrors and bandpass filtered. Timestamps of behavioral events were collected and read into the RZ5 system.

### Fiber Photometry Analysis

Signals from the 465 (GCaMP6) and 560 (tdTomato) streams were processed using custom written scripts in accordance with previously published methods (Jacobs & Moghaddam, 2020). Briefly, ΔF/F photometry signal was z-scored around each event (action, punishment, or reward delivery) to the 5 sec window prior that event. Z-score normalized ΔF/F were also utilized to compute cross correlation scores between PL and LOFC signals across ± 1 sec time lags for seek and take action epochs.

### Statistical Analysis

Data are presented as mean ± SEM. All statistical tests were done using GraphPad Prism (Version 9) and utilized an alpha of .05. Detailed information and results from statistical tests can be found in supplemental methods and Table 1, respectively.

## Results

### Rats learn to associate reward-directed actions with variable risk of harm

Expression of GCaMP6s and tdTomato was largely confined to the OFC and mPFC and optical fibers for recording were placed in the lOFC and PL subregions (Figure 1A). Animals were trained in a chained schedule of reinforcement that was split into blocks with varying risk of mild footshock depending on the phase of the experiment (Figure 1B). In each phase, neuronal activity was recorded during both seek and take actions (Figure 1C). These included a session where both actions were safe (Phase I), in the first session where risk of footshock was introduced during the seek action and a later session after rats had learned of risk of footshock (Phase II, see supplemental methods), and in two consecutive extinction sessions where risk of footshock was removed (Phase III).

**Figure 1:**
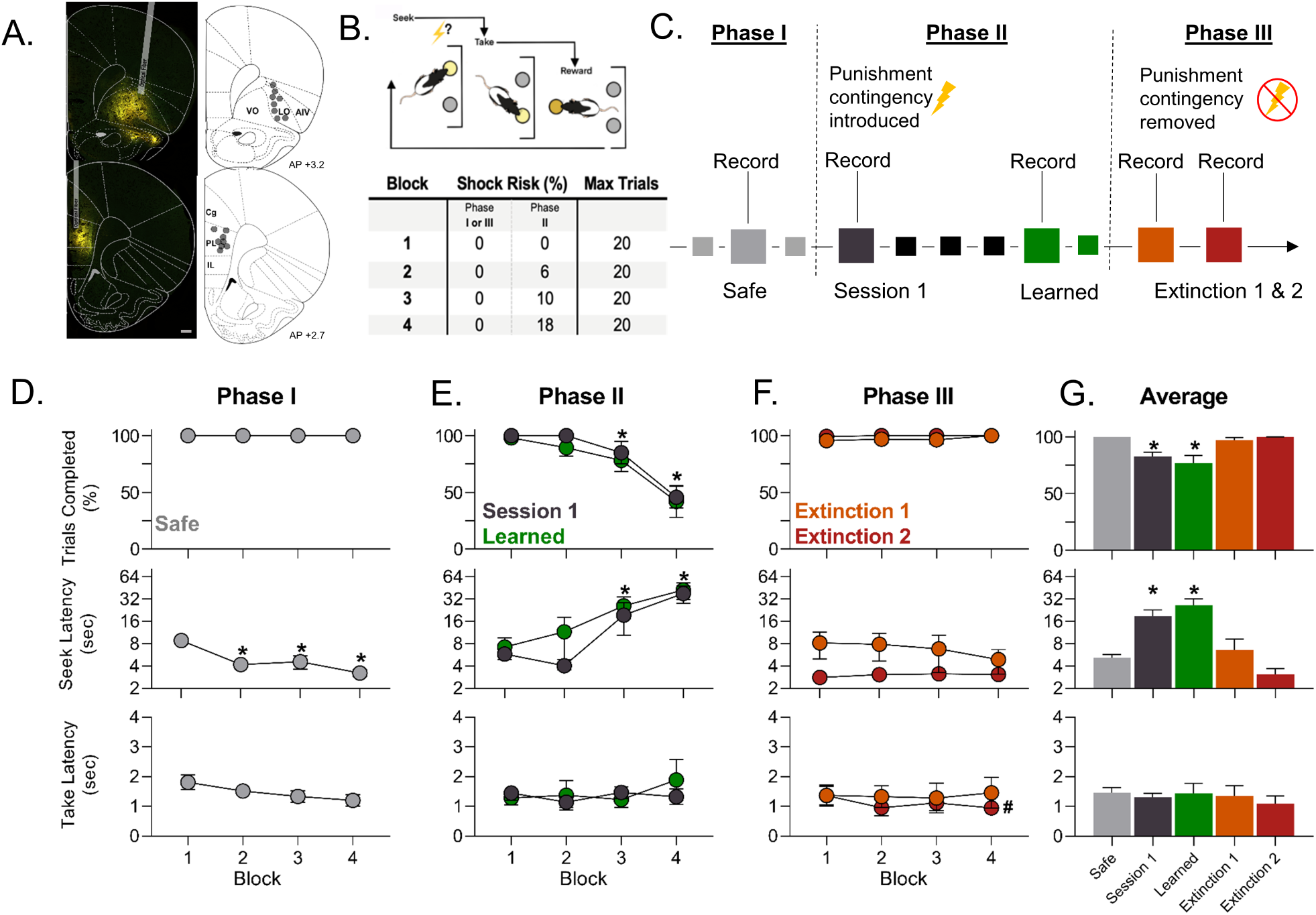
Experimental outline and behavior during each phase of the experiment. **A.** Expression of GCaMP and tdTomato was largely confined to the OFC and mPFC with fiber placements located in or just above lOFC (top) and PL (bottom) subregions. **B.** Task trial structure (Jacobs & Moghaddam, 2020) where successive seek and take actions must be performed to receive a reward and (bottom) structure of the session where the one hour session is split into 4 blocks, each with no risk of shock (Phase I or Phase III) or an ascending risk of footshock (Phase II). **C.** Out-line of fiber photometry recording sessions during each phase of the experiment. **D.** Behavior during Phase I when no risk of footshock was present. (Top) Trial completion over each block. (Middle) Latency to perform the seek action over each block. (Bottom) Latency to perform the take action over each block. *p*<*.05 vs. Block 1. **E.** Behavior during Phase II when animals learned of risk of footshock. (Top) Trial completion over each block. (Middle) Latency to perform the seek action over each block. (Bottom) Latency to perform the take action over each block. *p*<*.05 vs. Block 1 for combined Session 1 and Learned. **F.** Behavior during Phase III extinction training when animals learned risk of footshock was no longer present. (Top) Trial completion over each block. (Middle) Latency to perform the seek action over each block. (Bottom) Latency to perform the take action over each block. ^#^p*<*.05 vs Extinction 1. **G.** Average of trial completion, seek latency, or take latency for Blocks 2-4 for each phase of the experiment. (Top) Trial completion decreased in sessions with risk of footshock and recovered when risk of footshock was removed. (Middle) Latency to perform the seek action increased during Phase II sessions. (Bottom) Latency to perform the take action did not increase significantly over sessions. *p*<*.05 vs. Safe. n=5-8 rats in panel E, n= 8 rats otherwise. Scale bar= 500 µm. Data are presented as mean ± SEM.

Trial completion and action latencies for seek and take actions in each phase are depicted in Figure 1D-G. In Phase I, when all actions were safe, animals completed all trials for each block. Actions were performed rapidly for seek and take actions (Figure 1D) with seek action latencies becoming progressively faster within the session (*F_block_*(3,21)=18.1, p*<*.001). In Phase II sessions, when variable risk of footshock was present, rats showed decreases in trial completion during blocks 3 and 4 (*F_block_*(3,21)=31.04, p*<*.001; Figure 1E). These rats also showed large increases in latency to perform the seek action in the same blocks (i.e. the action associated with variable risk of footshock; (*F_block_*(3, 21) = 12.04, p*<*.001). Behavioral stability for Learned sessions was verified by a lack of significant effect of session when comparing the three consecutive sessions prior to the Learned recording session for trials completed and latency to execute either action (Supp. Figure 1A-C). Take action latency, which was always safe, did not increase over block in Phase II. In Phase III, when the risk of footshock was removed, rats adapted their behavior. Suppression of trial completion and increases in seek action latency dissipated in the first extinction session. By the second extinction session, all animals had returned to behavioral responses similar to that in Phase I (Figure 1F).

For analysis across phases, we collapsed data from the blocks with variable risk of footshock in Phase II (blocks 2-4) and compared them to the corresponding blocks in Phases I and III (Figure 1G). These findings demonstrated that both trial completion and seek action latency were impacted by the risk of punishment (F values *>* 9, p values *<*.001) as both Phase II sessions were significantly different from the Safe sessions (corrected p values *<* .04). Two additional post hoc tests also indicated higher seek latency in Phase II Learned sessions compared to both Phase III Extinction sessions (corrected p values *<* .01). Variable risk of footshock selectively affected the seek action, as the latency of the operationally similar take action was not changed in Phase II sessions (Figure 1G).

### lOFC and PL respond differently to variable risk of harm during reward seeking actions

In Phase I, lOFC response during seek action did not change over blocks (Supp. Figure 2, Table 1, Figure 2). The lOFC response to the take action did not change when collapsed over blocks but when activity was assessed by block, there was a negative response in block 1 compared to the neutral response in blocks 2-4 (*F_block_*(3, 21) = 5.69, p *<* .01, Figure 2B). The lOFC response in Phase I during reward delivery was not significant (p =.33 two-tailed; Supplement 2) and did not change over blocks (Figure 2C). In the PL, the seek action produced a phasic decrease in activity (p=.003 two-tailed; Supp. Figure 2) that persisted over all blocks (Figure 2D). In contrast, there was no significant phasic response during take action overall or across blocks (Figure 2E; Supp. Figure 2). PL reward response was positive throughout the session (p=.042 two-tailed; Supp. Figure 2) and did not vary with block (Figure 2F). Collectively, these findings characterize the basal state of population activity without threat of punishment and suggest minimal block related changes in action or reward encoding, with the exception of a transient decrease in lOFC response to the take action.

**Figure 2:**
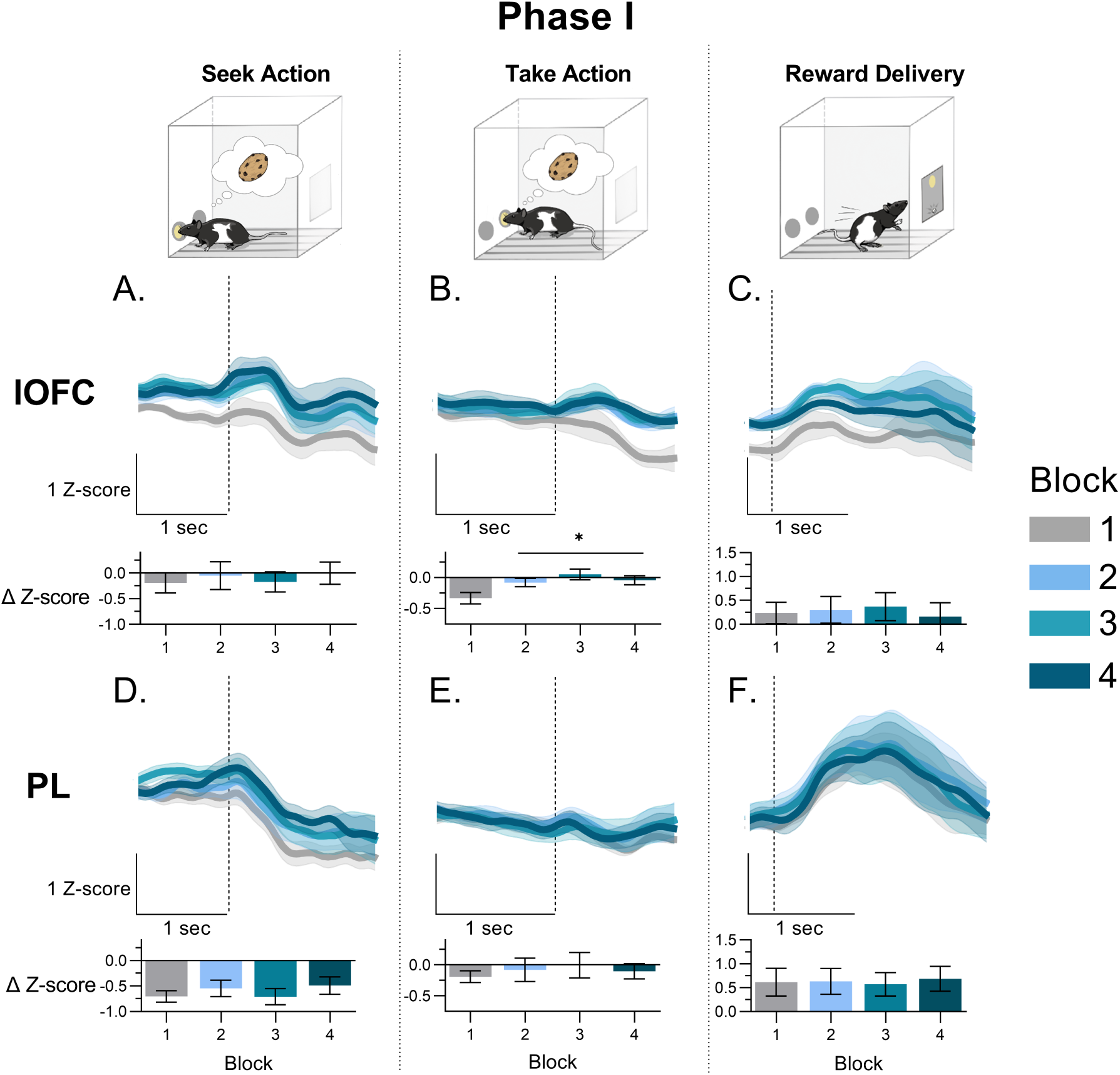
Action and reward responses in lOFC and PL during Safe sessions in Phase I. Onset of action or reward delivery is indicated by the dashed vertical line. **A.** Mean lOFC neuronal calcium activity during the seek action (top) and change scores for peri-action z-scored activity (bottom) over each of the four blocks. **B.** Mean lOFC neuronal calcium activity during the take action (top) and change scores for peri-action z-scored activity (bottom) over each of the four blocks. **C.** Mean lOFC neuronal calcium activity during reward delivery (top) and change scores for peri-reward z-scored activity (bottom) over each of the four blocks. **D.** Mean PL neuronal calcium activity during the seek action (top) and change scores for peri-action z-scored activity (bottom) over each of the four blocks. **E.** Mean PL neuronal calcium activity during the take action (top) and change scores for peri-action z-scored activity (bottom) over each of the four blocks. **F.** Mean PL neuronal calcium activity during reward delivery (top) and change scores for peri-reward z-scored activity (bottom) over each of the four blocks. * p*<*.05 vs. Block 1. Data are presented as mean ± SEM from n=8 rats. Vertical scale bars indicate 1 z-score. Horizontal scale bars indicate 1 second.

In Phase II, analyses focused on unpunished (Figure 3) and punished (Figure 4) trials separately due to the large increases in activity in the seek action period when concomitant with footshock (see below). In unpunished trials, lOFC activity during seek action did not change over blocks in Session 1 or after the task was learned (Figure 3A-B). In Session 1, there was no change in lOFC activity during take action (Figure 3C). After the task was learned, a phasic increase at take action execution emerged in lOFC in the last two blocks (*F_block_*(3, 18) =3.19, p=.048), though this failed to reach significance in Block 4 after post-hoc correction (Block 3 corrected p=.04, Block 4 corrected p=.057; Figure 3D). There was a weak positive response in the lOFC to reward delivery across blocks (Figure 3E-F). In the PL, the phasic decrease during seek action seen in Phase I was observed in Block 1 (where there was 0% risk of punishment) of Session 1. This pattern of activity was attenuated in the last block (18% risk) of Session 1 (corrected p value = .004; Figure 3G) and in the Learned session (corrected p value =.013; Figure 3H). This suggests that the PL is uniquely sensitive to actions with risk of punishment. This notion was supported by lack of significant change in the safe but operationally similar take action as a function of block in either session of Phase II (Figure 3I-J). Lastly, a phasic increase in response to reward was observed in the PL. Activity patterns were not different across blocks for reward delivery in both sessions indicating reward delivery responsiveness was not altered by variable risk of punishment (Figure 3K-L).

**Figure 3:**
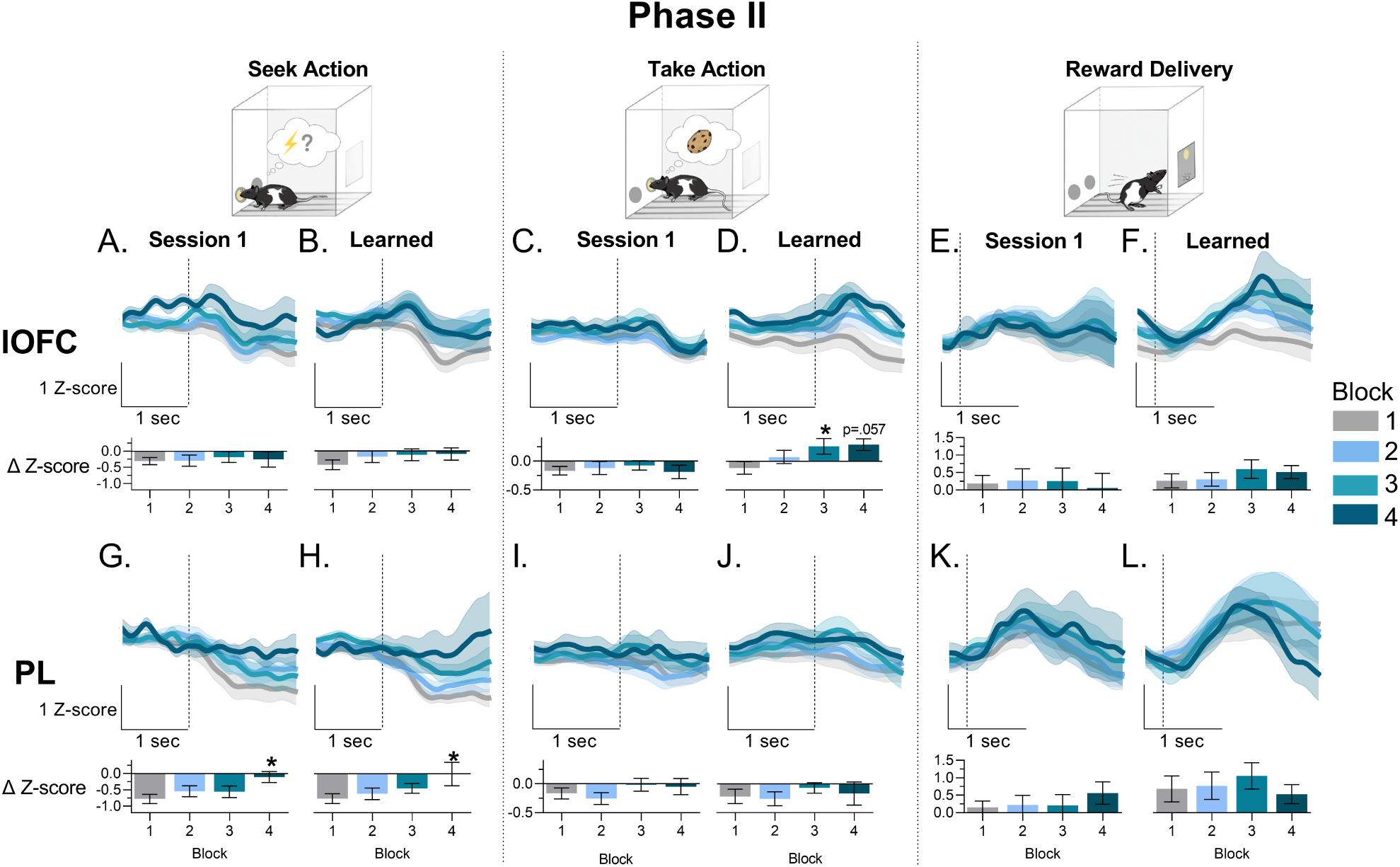
Action and reward responses in the lOFC and PL during probabilistic punishment sessions in Phase II. Onset of action or reward delivery is indicated by the dashed vertical line. Note: in these sessions each block was associated with an ascending risk of footshock for the seek action (0-18%). **A.** Mean lOFC neuronal calcium activity during the seek action (top) and change scores for peri-action z-scored activity (bottom) over each of the four blocks in Session 1. **B.** Mean lOFC neuronal calcium activity during the seek action (top) and change scores for peri-action z-scored activity (bottom) over each of the four blocks in Learned sessions. **C.** Mean lOFC neuronal calcium activity during the take action (top) and change scores for peri-action z-scored activity (bottom) over each of the four blocks in Session 1. **D.** Mean lOFC neuronal calcium activity during the take action (top) and change scores for peri-action z-scored activity (bottom) over each of the four blocks in Learned sessions. **E.** Mean lOFC neuronal calcium activity during reward delivery (top) and change scores for peri-reward z-scored activity (bottom) over each of the four blocks in Session 1. **F.** Mean lOFC neuronal calcium activity during reward delivery (top) and change scores for peri-reward z-scored activity (bottom) over each of the four blocks in Learned sessions. **G.** Mean PL neuronal calcium activity during the seek action (top) and change scores for peri-action z-scored activity (bottom) over each of the four blocks in Session 1. **H.** Mean PL neuronal calcium activity during the seek action (top) and change scores for peri-action z-scored activity (bottom) over each of the four blocks in Learned sessions. **I.** Mean PL neuronal calcium activity during the take action (top) and change scores for peri-action z-scored activity (bottom) over each of the four blocks in Session 1. **J.** Mean PL neuronal calcium activity during the take action (top) and change scores for peri-action z-scored activity (bottom) over each of the four blocks in Learned sessions. **K.** Mean PL neuronal calcium activity during reward delivery (top) and change scores for peri-reward z-scored activity (bottom) over each of the four blocks in Session 1. **L.** Mean PL neuronal calcium activity during reward delivery (top) and change scores for peri-reward z-scored activity (bottom) over each of the four blocks in Learned sessions. * p*<*.05 vs. Block 1. n=5-8 rats. Data are presented as mean ± SEM. Vertical scale bars indicate 1 z-score. Horizontal scale bars indicate 1 second.

**Figure 4:**
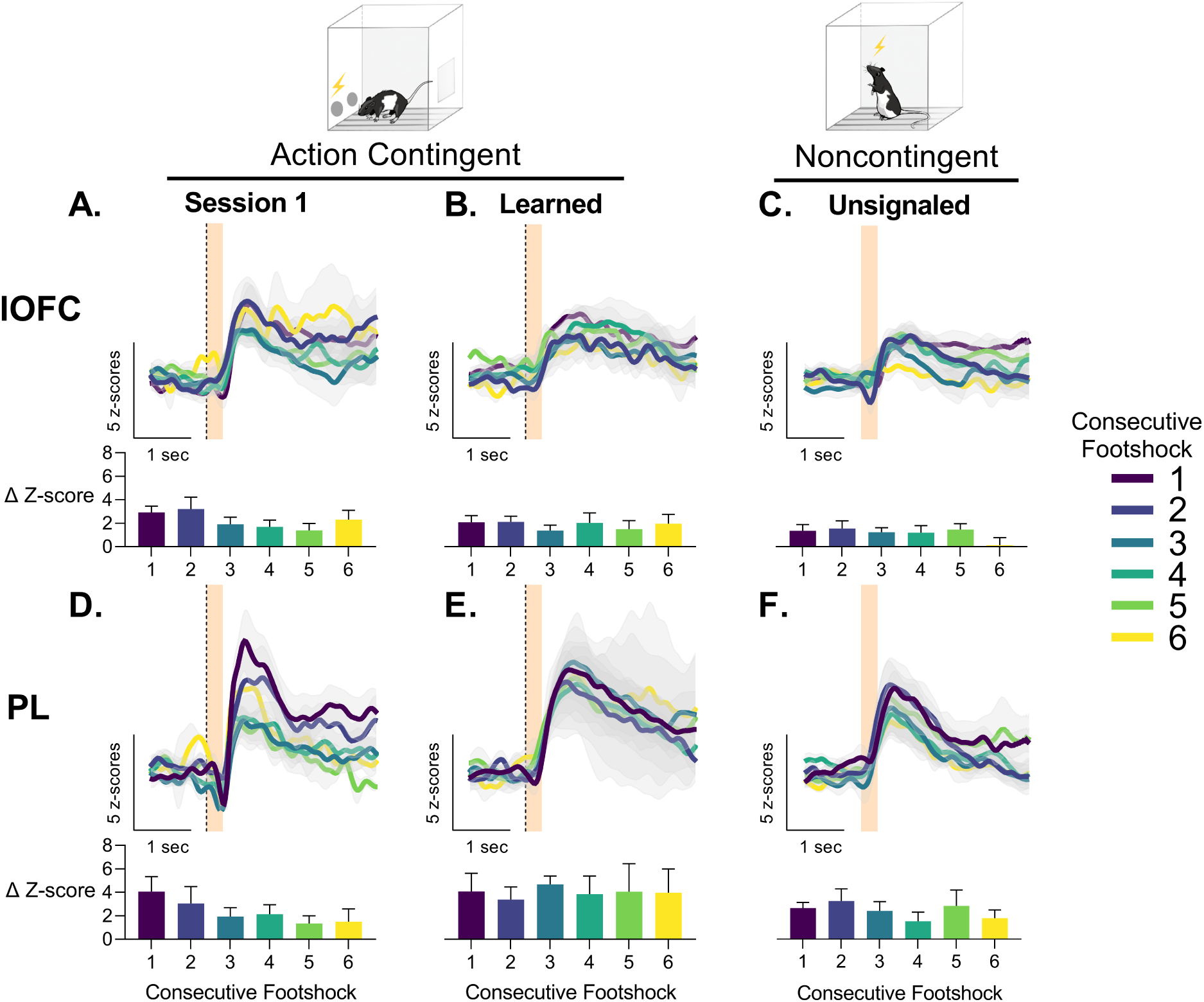
Response of the lOFC and PL at footshock onset (orange bar) over learning of variable risk of punishment in Phase II and when given noncontingently at the end of the experiment. Colors indicate the order of footshocks in that session. Dashed vertical lines indicate seek action (if applicable). **A.** Mean lOFC neuronal calcium activity following footshock (top) and corresponding change scores (bottom) in Session 1. **B.** Mean lOFC neuronal calcium activity following footshock (top) and corresponding change scores (bottom) in Learned sessions. **C.** Mean lOFC neuronal calcium activity following unsignaled footshock (top) and corresponding change scores (bottom) when given non-contingently. **D.** Mean PL neuronal calcium activity following footshock (top) and corresponding change scores (bottom) in Session 1. **E.** Mean PL neuronal calcium activity following footshock (top) and corresponding change scores (bottom) in Learned sessions. **F.** Mean PL neuronal calcium activity following unsignaled footshock (top) and corresponding change scores (bottom) when given non-contingently. n=2-8 rats. Data are presented as mean ± SEM. Vertical scale bars indicate 5 z-scores. Horizontal scalebars indicate 1 sec.

### lOFC and PL exhibit phasic response to action contingent and noncontingent footshock

Both lOFC and PL displayed a robust phasic increase to footshock when it was action contingent (one sample t values *>* 4.1, p-values *<* .005 averaged over all shocks). In lOFC no evidence of desensitization was observed as there was a phasic increase in neuronal response over all footshocks within Session 1 (Figure 4A) and in Learned sessions (Figure 4B) of Phase II. When these rats received the same footshock in a novel context noncontingently, the lOFC response to footshock was sustained and did not change significantly within the session (one sample t = 3.5, p = .0096; Figure 4C). In the PL, increases in neuronal calcium activity following action contingent footshock were observed in Session 1 and the Learned sessions. Though some trends towards a within session decrease were seen in Session 1, these were not statistically significant (*F_shock_*_#_(5,23)=.85, p=.53) and should be interpreted with caution as a small number of subjects (n=2) were willing to continue reward seeking after five foot-shocks. No within session changes in footshock response were observed in Learned sessions (Figure 4D-E). PL neuronal activity also did not desensitize in response to noncontingent footshocks (one sample t = 3.48, p = .01 averaged over all shocks; also see Figure 4F). These results indicate that contingent or noncontingent footshock stressors increase population level activity in lOFC and PL.

### Extinction of variable risk of harm results in rapid region-specific neural changes

To investigate neuronal adaptation in lOFC and PL when risk of harm was removed, we recorded from these regions during two consecutive extinction sessions (Phase III). In these sessions, reward contingencies were the same but punishment contingencies during seek actions were no longer present. Neuronal activity patterns in lOFC and PL differed during Phase III. In the lOFC, response to the seek action did not change over blocks and was characterized by a net neutral response in both extinction days (Figure 5A-B). Response to the take action, which had increased with risk of punishment in lOFC, no longer increased in the last block(s) in Extinction Day 1 or 2 (Figure 5C-D). Interestingly, the response to reward in the lOFC transiently increased over blocks (*F_Block_*(3, 21) = 6.91, p=.002). This was specific to blocks 2 and 3 of the Extinction Day 1 (corrected p values *<*.042, Figure 5E) but not Extinction Day 2 (Figure 5F). In the PL the phasic decrease in activity during seek action reemerged across all blocks similar to that of Phase I Safe sessions (Figure 5G-H). While seek action response changed in PL, the response to take action (Figure 5I-J) and reward delivery (Figure 5K-L) was similar to that observed over Phases I and II, and did not change as a function of block.

**Figure 5:**
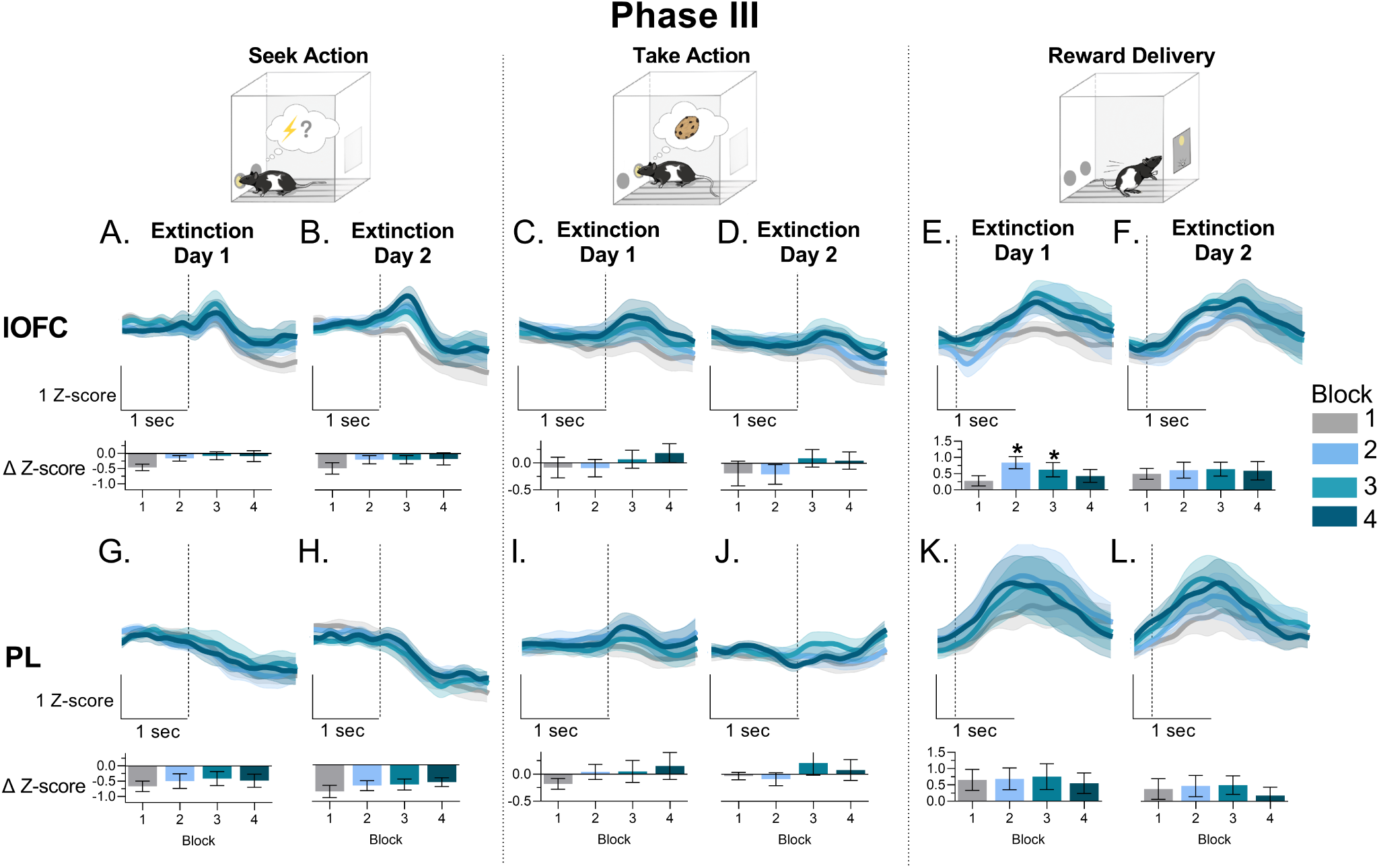
Response of the lOFC and PL during extinction sessions in Phase III. Onset of action or reward delivery is indicated by the dashed vertical line. Note in these sessions each block had no risk of footshock. **A.** Mean lOFC neuronal calcium activity during the seek action (top) and change scores for peri-action z-scored activity (bottom) over each of the four blocks in Extinction Day 1. **B.** Mean lOFC neuronal calcium activity during the seek action (top) and change scores for periaction z-scored activity (bottom) over each of the four blocks in Extinction Day 2. **C.** Mean lOFC neuronal calcium activity during the take action (top) and change scores for peri-action z-scored activity (bottom) over each of the four blocks in Extinction Day 1. **D.** Mean lOFC neuronal calcium activity during the take action (top) and change scores for peri-action z-scored activity (bottom) over each of the four blocks in Extinction Day 2. **E.** Mean lOFC neuronal calcium activity during reward delivery (top) and change scores for peri-reward z-scored activity (bottom) over each of the four blocks in Extinction Day 1. **F.** Mean lOFC neuronal calcium activity during reward delivery (top) and change scores for peri-reward z-scored activity (bottom) over each of the four blocks in Extinction Day 2. **G.** Mean PL neuronal calcium activity during the seek action (top) and change scores for peri-action z-scored activity (bottom) over each of the four blocks in Extinction Day 1. **H.** Mean PL neuronal calcium activity during the seek action (top) and change scores for peri-action z-scored activity (bottom) over each of the four blocks in Extinction Day 2. **I.** Mean PL neuronal calcium activity during the take action (top) and change scores for peri-action z-scored activity (bottom) over each of the four blocks in Extinction Day 1. **J.** Mean PL neuronal calcium activity during the take action (top) and change scores for peri-action z-scored activity (bottom) over each of the four blocks in Extinction Day 2. **K.** Mean PL neuronal calcium activity during reward delivery (top) and change scores for peri-reward z-scored activity (bottom) over each of the four blocks in Extinction Day 1. **L.** Mean PL neuronal calcium activity during reward delivery (top) and change scores for peri-reward z-scored activity (bottom) over each of the four blocks in Extinction Day 2. * p*<*.05 vs. Block 1. n=8 rats. Data are presented as mean ± SEM. Vertical scale bars indicate 1 z-score. Horizontal scale bars indicate 1 second.

To compare regions and sessions in the same statistical model, and account for any effects that could be due to individual variability in threat sensitivity, we collected data from the last block with at least 25% trial completion for each rat in Phase II sessions (i.e. when individual animals were still responding but demonstrated increased latency to complete risky actions). This corresponded to block 3 or 4 across all rats. We then compared changes in neuronal calcium activity in Phase II to the average of blocks 3-4 in Safe and Extinction sessions of Phases I and III. For footshock activity, we took the average of all footshocks over the sessions.

We observed a significant region by session interaction for the seek action (*F_int_*(4, 28) = 3.36, p =.023; Figure 6A-B). There was no change in the seek action over different Phases in the lOFC (p values *>*.49), but PL neural activity selectively changed in Session 1 and Learned sessions of Phase II (corrected p-values *<*.015, one-tailed) and returned to Phase I levels in Phase III. For the take response, we also observed an interaction between regions and session (*F_int_*(4, 28) = 4.878, p = .0041; Figure 6A-B). The take response in the lOFC was elevated in the Learned sessions compared to Safe sessions (corrected p value=.007, one tailed) whereas no change was observed between sessions in PL. While we had observed some changes in reward response for the lOFC (see Figure 5), there were no significant changes in response to reward delivery over sessions, between regions, nor interactions (F values *<* 1.17, p values *≥* .34). The footshock response revealed a main effect of region with PL response being greater than lOFC response (F(1,7) = 6.34, p = .039, Figure 6C), while the effect of Session failed to reach significance (F(2,14) = 3.18, p = .07). These findings highlight different patterns of activity across PFC subregions during the learning of presence and absence of variable risk of harm. Region-specific patterns of neuronal activity are schematized in Figure 6D. Specifically, we observed action specific adaptations to the safe action after threat is learned in the lOFC. In the PL, on the other hand, we observed an adaptation to the risky action associated with threat and a stronger response to the punishment itself while no changes in response to safe actions were observed. Importantly, these responses are highly flexible and dissipate when the risk of punishment is no longer present during threat extinction.

**Figure 6:**
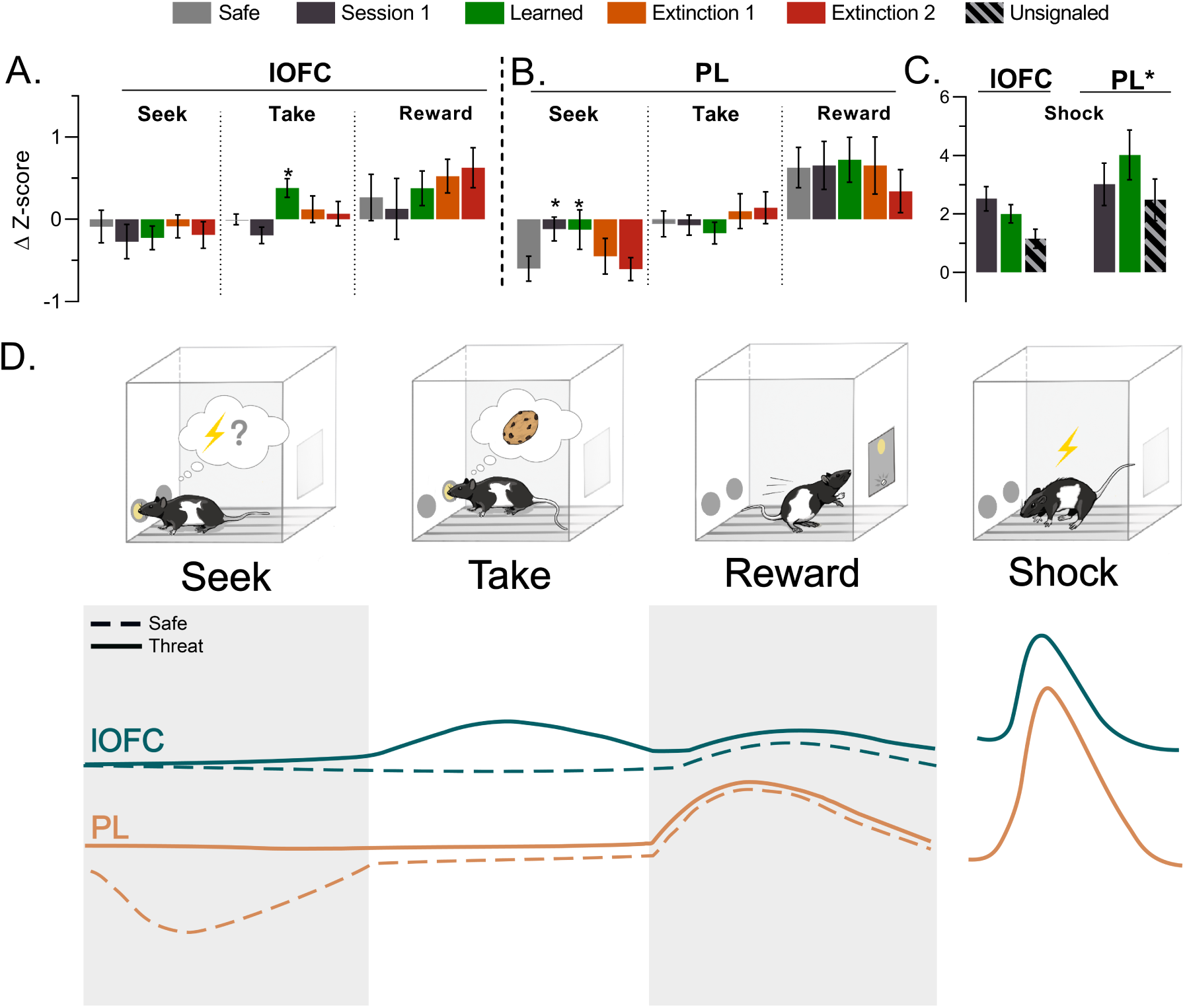
Peri-event population level neuronal calcium activity for lOFC and PL across all phases in the experiment (mean ± SEM) and summary schematic for main findings. Bar color indicates a given session. **A.** Peri-event change scores for seek, take, and reward in the lOFC. *p*<*.05 vs. Safe. **B.** Peri-event change scores for seek, take and reward in the PL. *p*<*.05 vs. Safe. **C.** Peri-event change scores for footshock in Phase II and unsignaled for PL and lOFC. *p*<*.05 PL vs. lOFC. **D.** Schematic of main findings for neuronal activity changes seen over the course of the experiment for each action and/or event. Solid lines indicate threat is present, while dashed lines indicate safe conditions.

### Correlated activity between frontal cortex subregions is reduced by variable risk of harm

PL and OFC are reciprocally and multi-synaptically connected. We, therefore, assessed correlated activity during seek and take actions as a proxy to infer synchronous activity across these regions that may reflect network dynamics (Sych et al., 2019). In Safe sessions (Phase I), we observed a positive correlation between PL and lOFC for both actions (Figure 7A). This positive correlation was also seen in the first three blocks of Session 1 when risk of harm was introduced but dissipated selectively for the seek action in Block 4 (Figure 7B). In the Learned sessions, positive correlations were seen in Block 1 and 2. In Blocks 3 and 4, however, the positive correlation was attenuated or blocked for the seek action (Figure 7C). In Phase III, the positively correlated activity during seek action reemerged Figure 7D-E). We observed a significant interaction between Session and Block, which was associated with suppressions of correlated activity in Phase II and III sessions (*F_int_*(12, 2644) = 6.54, p*<*.001). When comparing to Phase I, there was an attenuation of correlated activity for Block 4 of Session 1 and Blocks 3 and 4 in Learned sessions of Phase II (corrected p values *<*.01; Figure 7F). Furthermore, Block 4 of Extinction Day 1 remained attenuated (corrected p=.036; Figure 7F) but not Extinction Day 2 (corrected p=.246; Figure 7F). These analyses suggest that the correlated activity between PL and lOFC flexibly changes as animals learn to associate risk of harm with an action.

**Figure 7:**
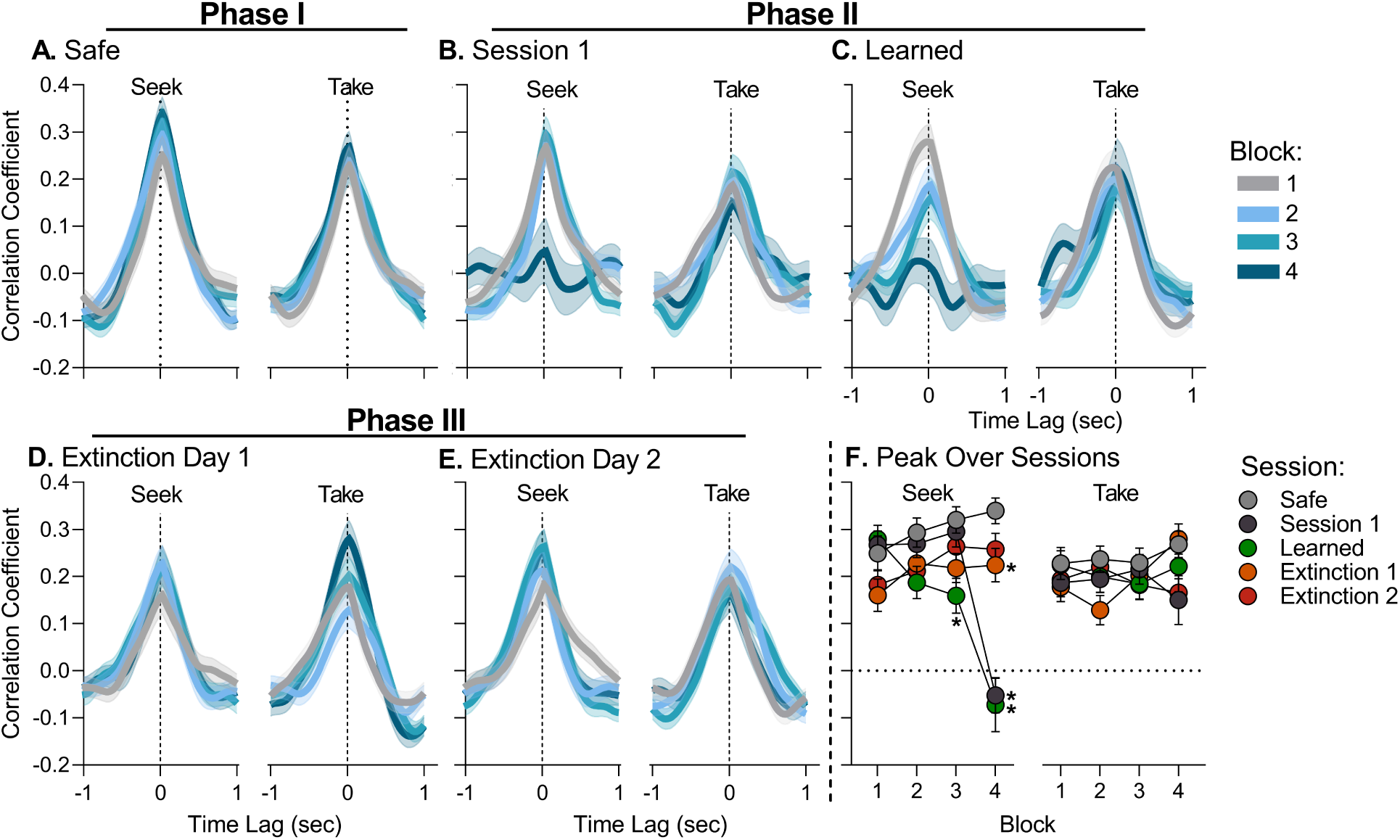
lOFC-PL cross correlation functions for each block (**A-E**) and the absolute peak correlation over each block of each session (**F**) for seek and take actions. Colors of lines indicate block and color of circles indicates session. **A.** Cross correlation for Safe sessions for seek (left) and take (right) actions over blocks. **B.** Cross correlation for Session 1 of Phase II for seek (left) and take (right) actions over blocks. **C.** Cross correlation for Learned sessions of Phase II for seek (left) and take (right) actions over blocks. **D.** Cross correlation for Extinction Day 1 of Phase III for seek (left) and take (right) actions over blocks. **E.** Cross correlation for Extinction Day 2 of Phase III for seek (left) and take (right) actions over blocks. **F.** Peak (absolute maximum) of the cross correlation function for each block of each session. Correlated activity in Block 4 was attenuated compared to Safe in Session 1, Learned, and Extinction 1. * p*<*.05 vs Safe session. n=51-160 trials from 5-8 rats. Data are presented as mean ± SEM.

## Discussion

Anxiety is a common and debilitating symptom of many brain disorders. Human imaging studies have consistently implicated PFC networks in anxiety (Balderston, Liu, et al., 2017; Bishop, 2009; Roberts, 2020; Sarlitto et al., 2018; Shin & Liberzon, 2010) including disruption of cognitive function by threat of harm in the clinical NPU task (Balderston, Vytal, et al., 2017). Preclinical research, typically focusing on innate anxiety models such as the elevated plus maze, has led to many insights into affective processes (Adhikari et al., 2010, 2011; Gunaydin et al., 2014; Loewke et al., 2021; Roberts, 2020). Anxiety in the real world, however, typically develops from the experience of learned associations between motivated actions and harmful outcomes (Grillon, 2008; Jacobs & Moghaddam, 2021; Park & Moghaddam, 2017a). The present experiments used a recently developed behavioral model for this mode of anxiety (Jacobs & Moghaddam, 2020) to assess the role of two cortical regions, lOFC and PL, in learning and expression of associating actions with a variable risk of harm. The model utilized two operationally similar actions, the first was associated with varying threat of punishment while the second was always safe and led to reward delivery. We found that PL neuronal activity selectively adapted to actions associated with threat, whereas lOFC neuronal activity adapted to safe actions. We also observe that correlated activity between the PL and lOFC was attenuated after learning of threat of harm, but was restored after threat was removed. These findings inform our understanding of the neuronal correlates of learned anxiety by demonstrating distinct roles for PL and lOFC in the representation of action during learning of anxiogenic contingencies.

### Divergent activity between frontal cortex subregions in learned anxiety

Frontal cortex subregions serve different roles in reward-motivated and emotional behaviors (Han et al., 2016; Holmes & Wellman, 2009; Hong et al., 2019; Izquierdo, 2017; Milad & Rauch, 2007; Roberts, 2020; Shiba et al., 2016; Simon et al., 2015; Stalnaker et al., 2015). Rodent studies using innate models of anxiety have found little effect of lOFC inactivation on anxiety-like behavior (Green et al., 2020; Lacroix et al., 2000; Orsini et al., 2015). In PL, innate anxiety studies have found PL neurons respond to transitions from safe to risky areas (Adhikari et al., 2011; Loewke et al., 2021), but inactivation studies have yielded mixed results (Bissiere et al., 2006; Kumar et al., 2013; Lacroix et al., 1998; Pati et al., 2018; Stern et al., 2010). In studies with threat during reward seeking, both lOFC and PL demonstrate increases in neural activity as indicated from c-Fos (Pascoli et al., 2015). Inactivation of the lOFC in contexts associated with threat result in hyper-suppression of action that permeates into trials with no threat (Orsini et al., 2015; Verharen et al., 2019). PL inactivation, however, enhances action under threat and disrupts adaptation to varying risk of punishment (Chen et al., 2013; Orsini et al., 2018; Resstel et al., 2008; Verharen et al., 2019). Together these findings suggest roles for the lOFC and PL in the discrimination of, or adaptation to, safety and threat, respectively. These studies justify the current focus on lOFC and PL in learning of action-threat contingencies.

We have recently reported on a novel approach to model anxiety in rodents where reward-guided actions are associated with a risk of punishment (Jacobs & Moghaddam, 2020; Park & Moghaddam, 2017b). This paradigm is distinct from typical preclinical approaches because it models anxiety in the context of learning that actions may be associated with a variable threat of harm. Using this approach, we find that neuronal activity in lOFC and PL differentially changed as a function of learned anxiety. In lOFC an increase in activity was observed for safe actions, after rats learned of the presence of threat. This was distinct from PL activity, which showed no learning or threat related changes following the safe action. Instead PL activity adapted to risky actions when threat was present. While insight into the subtypes of neurons that reflect these changes is outside the scope of the current studies, these findings suggest that changes in population activity state during action execution are different in these two PFC subregions in response to anxiety.

Given that pathological anxiety is partially characterized by heightened risk assessment in the absence of threat, we also determined how these neuronal correlates of learned anxiety changed as rats learned that there was no longer a threat of harm. These studies found concomitant changes in lOFC and PL neuronal activity as learned action-threat associations were extinguished. Neuronal activity changes were again action specific, with both lOFC and PL returning to baseline activity patterns for safe and risky actions, respectively.

Punishment itself increased lOFC and PL activity regardless of exposure history or whether punishment was action contingent. This finding corroborates very recent longitudinal experiments which reported stable PL response to footshock (though drift in individual cells was noted) and provide insight into the paucity of data characterizing lOFC response to footshock stressors (Kim et al., 2017; Patel et al., 2022; Turner et al., 2021). Our design also allowed us to directly compare this response in each region within subjects. In this regard, footshock related neuronal activity was greater in the PL compared to lOFC, which may indicate a greater role for PL in response to stressors and threats.

### Correlated activity between lOFC and PL is disrupted by learned anxiety

Threat of punishment disrupted correlated activity between the lOFC and PL during risky action. This finding is consistent with human studies showing anxiety disrupts network activity in several brain regions including frontal cortex (Balderston, Vytal, et al., 2017; Cornwell et al., 2017; Sartori & Singewald, 2019). Disruptions in correlated activity fully recovered after two days of extinction training, when neuronal and behavioral responses returned to baseline. This finding is in line with studies in humans that found symptom improvement after emotional regulation therapy in GAD patients correlates with changes in functional connectivity with frontal cortex (Scult, Fresco, et al., 2019), and complements human and non-human primate studies that suggest the regulation of anxiety involves intact cortical connectivity with other brain regions (Kenwood, Oler, et al., 2022; Scult, Knodt, et al., 2019).

In rats, correlated activity between lOFC and mPFC can form a common value signal following reward (Amarante & Laubach, 2021). These value signals are believed to be critical in supporting learning in reward-driven tasks (Miller et al., 2022). Thus our findings may represent a mechanism for how anxiety impacts neural computations to support adaptive behavior (Hartley & Phelps, 2012; Park et al., 2016). These results add to previous work in rats where mPFC-VTA correlated activity was attenuated during risky action (Jacobs et al., 2022; Park & Moghaddam, 2017b) and highlight the involvement of multiple neural pathways in the effects of anxiety on motivated behavior.

### Implication of our findings for theories related to approach-avoid decision making

Using reinforcement sensitivity theory as a framework for approach-avoid decision processes (Gray, 1984; McNaughton & Corr, 2004), our findings of discrete adaptive response to safe versus risky actions place PFC subregions into distinct systems. Reinforcement sensitivity theory posits three systems which underlie approach-avoidance conflict: the behavioral activation system (BAS) for approach, fight-flight-freeze system (FFFS) for avoidance, and behavioral inhibition system (BIS) which promotes vigilance and risk assessment when engaged due to sufficient co-activation of BAS and FFFS (see McNaughton & Corr, 2004; McNaughton & Gray, 2000). While the frontal cortex is suggested to be involved in many of these systems, its role has been described as “tentative” (Corr, 2013).

The observed changes in lOFC representation of the safe action after learning of threat may indicate lOFC involvement in the BAS, where lOFC may support enhanced BAS drive to temper the inhibitory systems when particular actions are safe. Our findings of changes in PL representation of threat associated actions and larger overall response to punishment may place the PL in the BIS, where threat assessment and attention allocation are core functions. These interpretations are in line with prior work. lOFC lesion in contexts with threat risk results in a decrease in approach behavior, even when risk is at 0% (Orsini et al., 2015; Ray et al., 2018; Shiba et al., 2016). PL inactivation, however, has been shown to result in compulsive reward seeking under conflict (Chen et al., 2013; Friedman et al., 2015), suboptimal adaptation to changes in threat probability (Orsini et al., 2018), as well as disrupted attention to threatening stimuli during fear learning (Sharpe & Killcross, 2015, 2018); which are consistent with PL as a hub for attention to threat.

These findings also provide a basis for why both regions are commonly implicated in maladaptive approach and avoidance behaviors. Without proper engagement of the BIS during learning of threatening contingencies via PL, proper attention and deliberation processes would be weakened. This could produce maladaptive approach or avoidance as behavioral control would too rely on the BAS and FFFS in the absence of BIS mediation. Alternatively, if the BAS is not engaged when threat is distant via lOFC then maladaptive levels of suppression may emerge from unbridled fight-flight-freeze system activity.

## Conclusion

Anxiety in the real world may manifest after organisms learn that goal-motivated actions are associated with a risk of harm. In this regard, anxiety is not a standalone construct, but an ongoing modulator of decisions to approach or avoid. Our data address a void in preclinical data for this mode of anxiety and find that two frontal cortex subregions have complementary roles during the learning and extinction of learned anxiety. Collectively our results suggest that distinct adaptation to safe and threatening actions in, and disruption of correlated activity between, these two regions may be a mechanism by which anxiety can influence frontal cortex processes. These findings inform the clinical literature and our overall understanding of how overlapping cortical systems are implicated in reward-guided action execution during anxiety.

## Supplemental Methods

Return to main methods

### Subjects

Adult (n=8) Long-Evans rats, housed on a reverse 12 h:12 h light/dark cycle, were used. Animals were pair housed until implantation of the optical fiber. All experimental procedures and behavioral testing were performed during the rodents’ dark (active) cycle. Both males (n=6) and females (n=2) were utilized. Subjects were run in several cohorts with both males and females insomuch as possible and were bred in house.

### Surgery

#### Viral infusion Surgery

Subjects were injected with AAV8-hSyn-GCaMP6s-P2A-tdTomato (OHSU Vector Core, 5e13 ng/mL) to allow for pan-neuronal expression of fluorescent calcium indicator GCaMP6s and the non-calcium dependent fluorophore tdTomato. TdTomato has been used by us and others as a motion artifact for fiber photometry recording in rodents (Babayan et al., 2018; Jacobs & Moghaddam, 2020; Matias et al., 2017; Menegas et al., 2018). Rats were anesthetized via isoflurane and placed in a stereotaxic apparatus. Following an incision and topical application of lidocaine, craniotomy was performed to lower a 10 uL microsyringe (Hamilton) to infuse virus into the PL and lOFC. Two injections were made (325 nL/site @ 50 nL/min) at the coordinates AP + 2.8, ML ± 0.65, DV −2.5 and −3.5 mm (from dura) and AP + 3.0, ML ± 3.0, DV −3.8 and −4.8 mm (from dura), for the PL and lOFC respectively, with the most ventral injection always performed first. A microcontroller (World Precision Instruments) controlled the amount and rate of the injections. Virus was allowed to diffuse for 5 minutes after the most ventral injection. The needle was then slowly raised and the second injection was performed and allowed 12 minutes to diffuse. After this the needle was removed, the incision was stapled, and animals were given a 5 mg/kg injection of carpofen subcutaneously.

#### Fiber Implant Surgery

After allowing at least four weeks for virus expression, subjects were implanted with an optical fiber aimed at the PL (AP +2.8, ML ± 0.7, DV −3.3 mm from dura) and lOFC (AP +3.0, ML ± 3.6, DV −5.2 mm from skull at a 7° angle) using the surgical procedures outlined above, with the exception that three additional bore holes were made for three skull screws which surrounded the craniotomy. The optical fiber was lowered and glued to the skull with light-curing epoxy (Tetric N-flow, Ivoclar Vivadent). Subjects were given 5 mg/kg of carpofen and returned to ad libitum food for 5 days before returning to food restriction. Subjects were given 1 week to recover from surgery before behavioral testing.

### Apparatus

An operant chamber was used for behavioral testing (Coulbourn Instruments, PA). The chamber included two nosepoke holes on one wall located 2 cm above a grid floor. The grid floor was connected to a shock generator which delivered footshocks. On the opposite wall was a food trough which dispensed sucrose pellets (45-mg, Bio-Serv #F0023) and detected food trough entries. The operant chamber had an opening in the top of the box to permit entry for the recording patchcord. For the unsignaled footshock sessions the same basic chamber was utilized but nosepoke holes and feeder troughs were replaced by metal squares (i.e. no operandi were present). Experimental stimuli were controlled through Graphic State software (version 4, Coulbourne Instruments) running on a windows PC.

### Chain Schedule Trianing

As previously described (Jacobs & Moghaddam, 2020), subjects were first trained to make two spatiotemporally distinct successive instrumental responses to receive a 45-mg sugar pellet under a fixed ratio one schedule of reinforcement (FR1). Chain schedule training began after habituation to the operant box and food trough (one session with a pellet dispensed on a VI-45 sec schedule). The first action will be referred to as the “seek” action and the second action will be referred to as the “take” action. Subjects were first trained to respond on the “take” nosepoke under a fixed ratio 1 (FR1) schedule. Sessions lasted until 80 pellets were delivered or 60 min elapsed. After this, subjects were trained to respond on the “seek” nose-poke. After the seek nosepoke was illuminated, a response on the seeknose poke (first link of the chain) resulted in extinguishing of the seek nosepoke light concurrent with a 1.5 sec delay. The take nosepoke was then illuminated and completion of a FR1 on the take nose poke (second link of the chain) turned off the take nosepoke light and, following a 1-sec delay, initiated pellet delivery and food trough illumination. Subjects were required to retrieve the pellet to initiate a 15 s intertrial interval (ITI). After the ITI, the seek nose poke was illuminated and a new trial began. The side of seek and take nosepokes were counterbalanced across subjects. After the completion of 80 trials or 60 min, the session was terminated. After this initial training period subjects were given implantation surgery for optical fibers (see above) and allowed at least one week to recover. Chained schedule behavior was then reacquired before subjects proceeded to Phase I of the experiment.

### Phase I - Safe Sessions

After chain-schedule training subjects moved to Phase I where they performed the same chained schedule of reinforcement but sessions were split into four 12 min blocks separated by a 2 min inter-block interval where all lights were extinguished. Subjects could complete up to 20 trials in each block (i.e. earn 20 reinforcers). In Phase I sessions, no risk of footshock was present for any action and subjects were given at least four Phase I sessions before beginning the probabilistic punishment task in Phase II and neuronal activity was recorded with fiber photometry after 3-5 sessions of training with these safe contingencies.

### Phase II Sessions - Probabilistic Punishment Task

The Probabilistic Punishment Task follows previously published methods (Jacobs & Moghaddam, 2020). Identical to the Safe sessions, Probabilistic Punishment Task sessions consisted of four blocks of 20 trials each. The take action-reward contingency remained constant. However, the probability of receiving a punishment (300 ms electrical footshock at 0.25 mA) after the seek action increased over the blocks (0%, 6%, 10%, and 18% in blocks 1, 2, 3, and 4, respectively). Ascending punishment risk was used to prevent overgeneralizing of the shock and is used in other risky decision procedures (Jacobs & Moghaddam, 2020; Park & Moghaddam, 2017b; St Onge & Floresco, 2010). Successful seek action execution always activated the take cue light. Neuronal activity was recorded in the first Probabilistic Punishment Task session and subjects were required to complete at least four Probabilistic Punishment Task sessions and show 3 consecutive sessions within 30% of the mean for the number of completed trials before neuronal activity was recorded again for the ‘Learned’ session. If subjects were completely resistant to footshock, we increased shock intensity (up to 0.5 mA) to achieve suppression of behavior for at least 3 days and neuronal activity was re-recorded for the ‘Learned’ session. This was required for two male rats. All subjects were given at least one week of Phase II (*≥* 7 sessions).

### Phase III - Extinction Sessions

Following learning of the Probabilistic Punishment Task, two extinction sessions were given by removing the probabilistic punishment contingency (i.e. all actions were safe) and keeping all other contingencies. Neuronal activity was recorded in both extinction sessions.

### Unsignaled Noncontingent Footshock Session

To determine if any learning related changes in footshock responsivity were products of foot-shock exposure. We applied the same shock (0.25 mA, 300 msec) in a different context (operant box with no nosepokes or feeder) with a variable 90 sec (60-120 sec) inter-stimulus interval. Subjects were given one session to acclimate to the new context for 10-min. In the next session subjects received six footshocks. Footshocks were administered after a 5-min habituation period and were not signaled. Unsignaled footshock sessions were performed after extinction.

### Fiber Photometry Apparatus

Recordings were performed with a commercially available fiber photometry system, Tucker-Davis Technologies RZ5. Recording was accomplished by providing both 470 nm and 560 nm excitation light through the 400-um core patchcord to the PL or lOFC for GCaMP6 and td-Tomato signals, respectively. LEDs were sinusoidally modulated at 210 and 330 Hz. Light intensity was initially set at 10 µW but adjusted for each subject to prevent photodetector clipping. This resulted in light levels of 1-12 µW. Emissions from GCaMP and tdTomato (500-540 nm and 580-680 nm, respectively) were collected back through the patchcord to dichroic mirrors and bandpass filters within a dual florescence Doric minicube. Fluorescence was converted to voltage through four femtowatt detectors (Newport 2151). Data were recorded using Synapse software (Tucker-Davis Technologies) and timestamps of behavioral events were collected by 5V TTL pulses that were read into the RZ5 system. Synapse software demodulated fluorescence signals in real time at 1 kHz with a 6-Hz low pass filter.

### Fiber Photometry Data Analysis

#### Peri-event analysis

Signals from the 465 (GCaMP6) and 560 (tdTomato) streams were processed in Python (Version 3.7.4) using custom written scripts in accordance with previously published methods (Jacobs & Moghaddam, 2020). Briefly, 465 and 560 streams were low pass filtered at 3 Hz using a butterworth filter and subsequently broken up based on the start and end of a given trial. The 560 nm signal was fitted to the 465 using a least-squares first order polynomial and subtracted from 465 signal to yield the change in fluorescent activity. The corrected signal was then z-scored for a given epoch (seek, take, reward delivery). The z-score comparison window was 5 sec ΔF/F prior to a given epoch to quantify changes in post action or outcome activity. Because of the magnitude of footshock response immediately following seek action and because we were interested in footshock response itself, we separated punished (footshock) and unpunished trials for plotting and analysis. To gain point estimates of peri-event responses we took the change score of mean z-scored activity. Windows were defined as the 1.5 or 1 seconds after a seek or take action, respectively and the 0.5 seconds before action. For reward delivery windows were defined as 1.5 second following delivery and compared to the timepoint immediately before delivery.

#### Time Lagged Cross-Correlation Analysis

Using z-score normalized ΔF/F, we computed covariance scores by taking the dot product of PL and lOFC signals across all possible time lags for +/- 1 second of seek and take action epochs. These covariance scores were normalized for each time lag to achieve a correlation coefficient between −1 and 1 using the following equation:

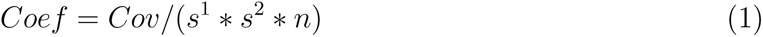

Where *Cov* is the covariance from the dot product of the signal for each timepoint, *s^1^* and *s^2^* are the standard deviation of the PL and lOFC streams, respectively, and *n* is the number of samples. An entire cross-correlations function was derived for each trial.

### Statistical Analysis

Detailed results of statistical tests are found in Table 1.

#### PPT Behavior

We analyzed trial completion and action latencies for seek and take action in accordance with our previous study (Jacobs & Moghaddam, 2020). Briefly, latency data were only included for blocks with at least two trials to prevent errant data points from skewing the data. For the take latency we only included unpunished trials to rule out changes in take action latency that were related to footshock receipt rather than hesitation to perform the action. Statistical procedures utilized repeated measures ANOVA or mixed-effects models using factors of risk block and/or session. Post-hoc comparisons were performed using Dunnett tests when comparing to Safe sessions or Block 1. Bonferroni correction was used if additional tests were added. Behavior data files were processed using custom written scripts in Python (version 3.0) and all statistical analyses were performed in GraphPad Prism (Version 8, San Diego, CA) and utilized an *α* of .05. All tests were two-tailed unless motivated by a directional hypothesis and are specified in Table 1.

#### Fiber Photometry

To assess peri-event response we took change scores for each block for each subject. Initially for the Safe session, change scores averaged over all blocks were compared by one sample t-test (comparing to 0) to see if any phasic response was observed at baseline. If we observed a violation in normality from the Shapiro-Wilk test, then a Wilcoxon signed rank test was used. To investigate effects of time on task (safe sessions) or risk of punishment (Phase II sessions) we then utilized separate one way ANOVA or mixed effects models with block as a factor for each session. Post hoc comparisons were corrected with Dunnett tests comparing to the first block. Lastly, to cross compare regions and sessions in the same model we utilized a two-way ANOVA with factors region and session for the final 3-4 blocks of each session. Post hoc comparisons were corrected with Dunnett tests comparing to the Safe session in Phase I. Tests were done using GraphPad Prism (Version 8) and utilized an *α* of .05. All tests were two-tailed unless motivated by a directional hypothesis and are specified in Table 1.

To assess correlated activity changes as a function of risk or session, we took the absolute max correlation and standard error for the overall cross correlation function. These values were compared by a two-way ANOVA with factors risk and session followed by post-hoc tests with a Dunnett correction. Tests were done using GraphPad Prism (Version 8) and utilized an *α* of .05.

### Excluded Data

Behavioral data points outside of 3 standard deviations from the mean were excluded as outliers which resulted in four data points being removed. Trials where the optical fiber patch cord fell off during action periods or needed to be reconnected were also excluded. Any trial with a z-score value *>* 40 was excluded as noise.

### Histology & Imaging

Viral expression and fiber placements were verified after behavioral studies. Subjects were transcardially perfused with 0.01 M phosphate buffered saline (PBS) and 4% paraformaldehyde (PFA). Brains were extracted and post-fixated in 4% PFA for 24 h before being placed in 20% sucrose solution and stored at 4°C. Forty-µm brain slices were collected on a cryostat (Leica Microsystems) and preserved in 0.05% phosphate buffered azide. Brain slices were mounted to slides and cover slipped with Vectashield anti-fade mounting medium (Vector Labs). A Zeiss Axio Observer microscope was used to image brain slices for GFP (Zeiss Filter set 38: 470-nm excitation/525-nm emission) and tdTomato (Zeiss Filter Set 43: 545-nm excitation/605-nm emission) to validate expression of both fluorophores in cells near the fiber tip. Fiber placement was determined by the brain slice demonstrating the most ventral fiber location.

Immunohistochemistry with a GFP antibody was used if a subject lacked virus expression to confirm the presence or absence of GCaMP6s. Brain slices were permeabilized in 3% BSA, 0.5% Triton X, and 5% Tween 80 dissolved in PBS + 0.0% sodium azide for 2 hr at room temperature. Slices were then incubated with mouse antiserum against GFP (Abcam, Catalogue# 1218, 1:500) diluted in PBS + Azide, 3% BSA + 0.1% Triton, and 1% Tween for 48 hr at 4°C. Slices were then washed in PBS + Azide, 3% BSA + 0.1% Triton + 1% Tween, three times for five minutes each. After this, slices were incubated with donkey-anti-mouse Alexa-488 (Abcam, Catalogue# 150105, 1:2000) diluted in PBS + Azide, 3% BSA + 0.1% Triton, and 1% Tween for 24 hr at 4°C and subsequently washed again as outlined above. Slices were then mounted to slides with Vectashield and imaged using the same procedures outlined above.

## Supplemental Tables & Figures

**Supplemental Data: Table 1.**
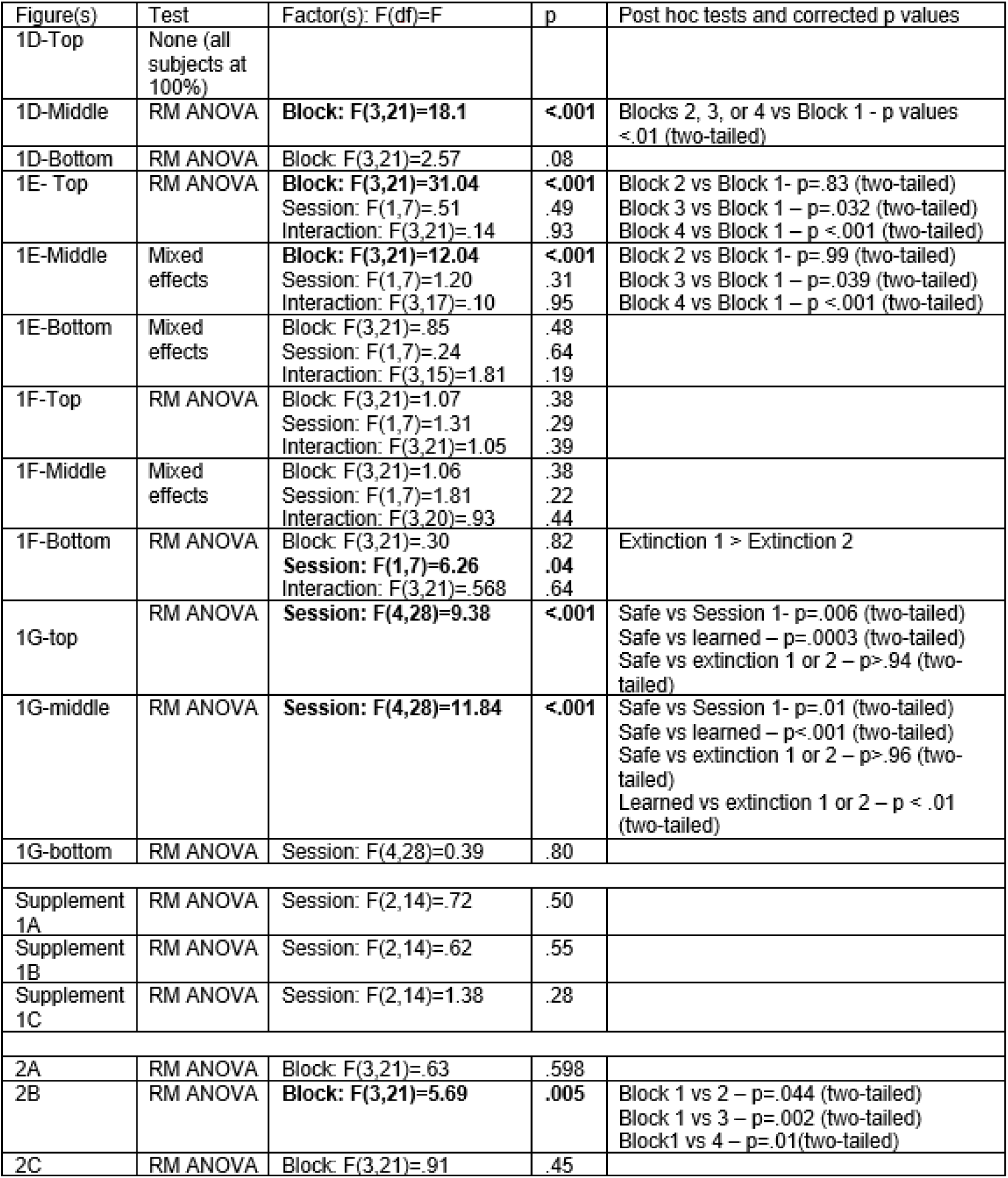
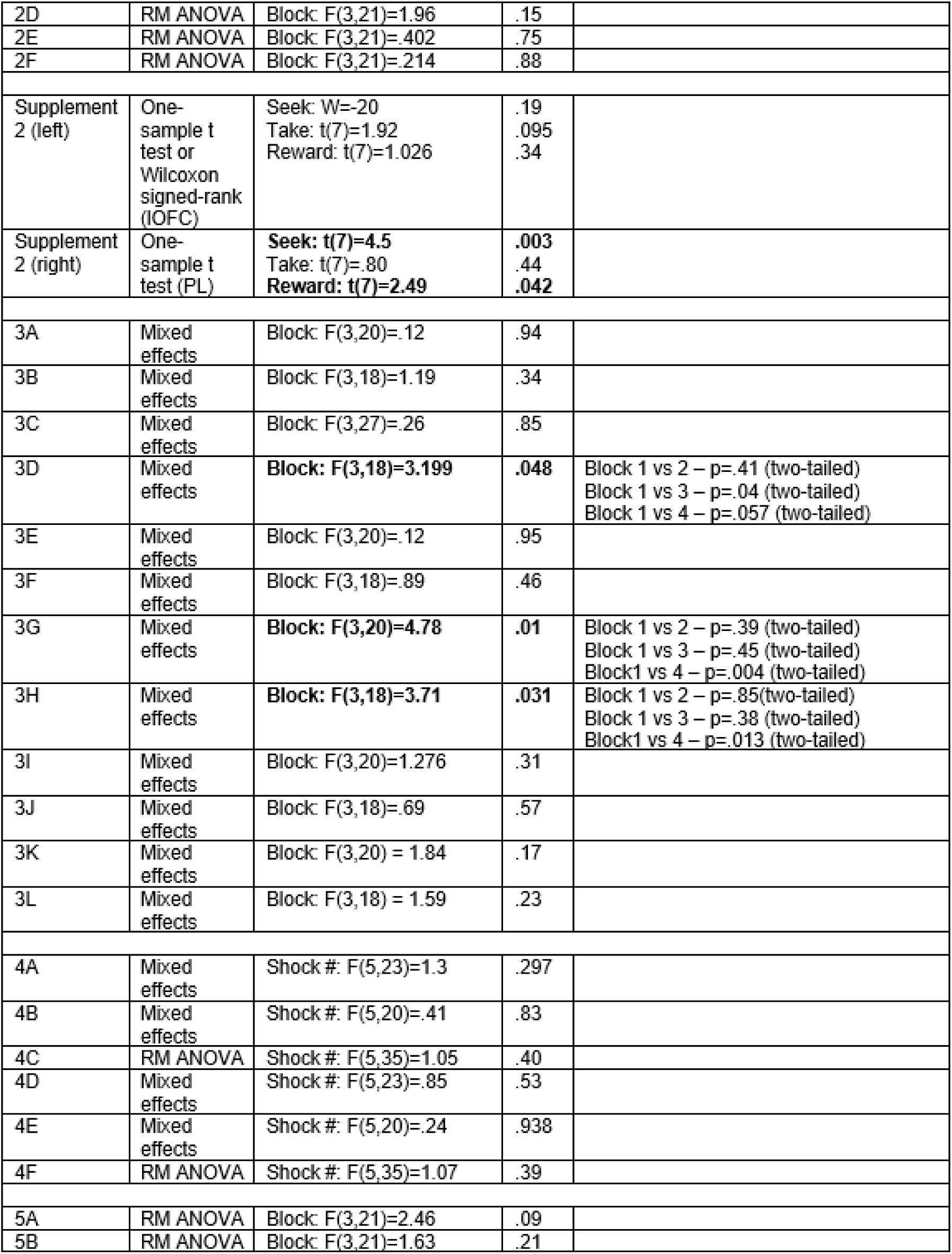
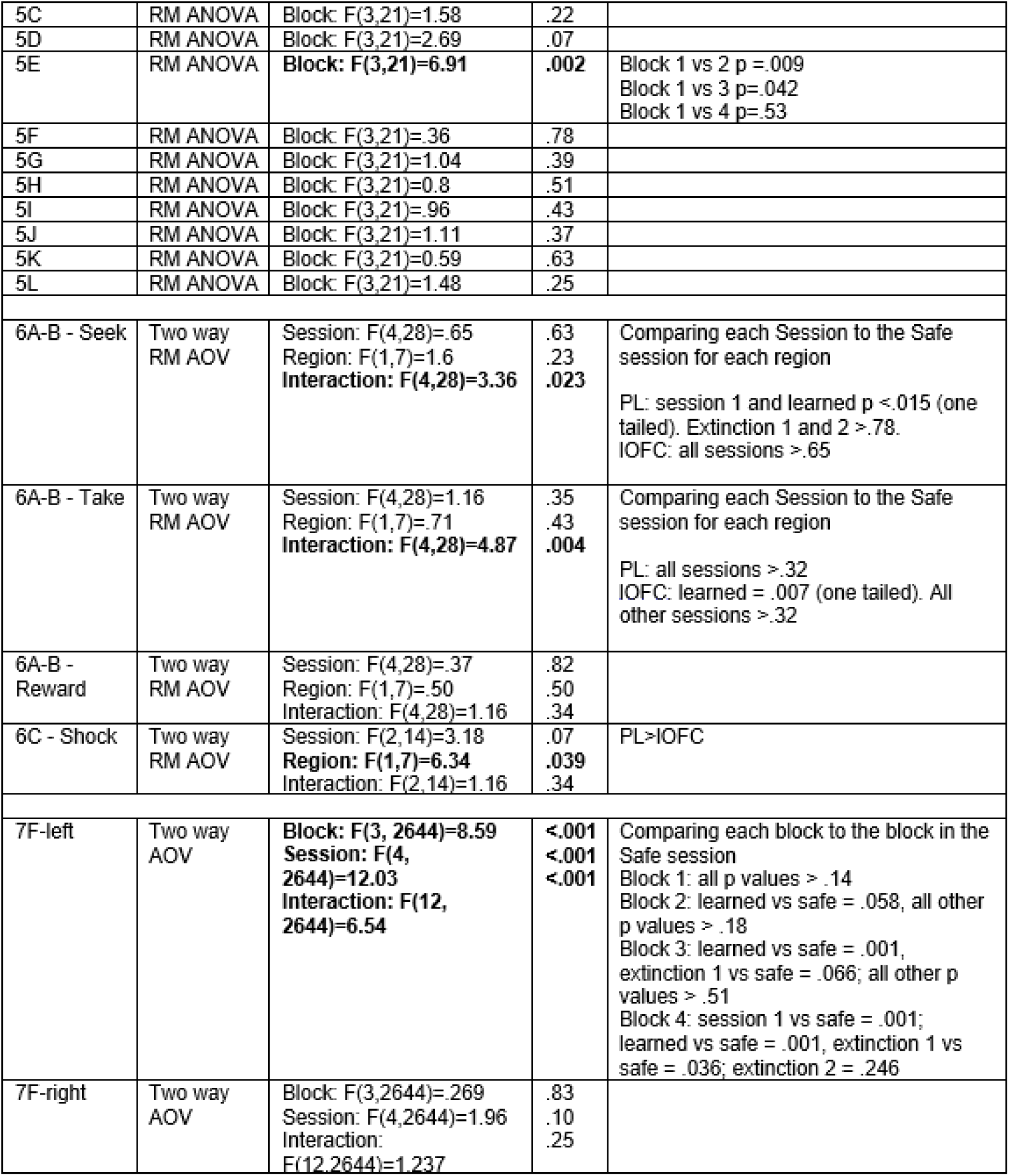
Results of Statistical Analyses.

| Figure(s) | Test | Factor(s): F(df)=F | p | Post hoc tests and corrected p values |
| --- | --- | --- | --- | --- |
| 1D-Top | None (all subjects at 100%) |  |  |  |
| 1D-Middle | RM ANOVA | **Block: F(3,21)=18.1** | **<.001** | Blocks 2, 3, or 4 vs Block 1 - p values <.01 (two-tailed) |
| 1D-Bottom | RM ANOVA | Block: F(3,21)=2.57 | .08 |  |
| 1E- Top | RM ANOVA | **Block: F(3,21)=31.04**<br>Session: F(1,7)=.51<br>Interaction: F(3,21)=.14 | **<.001**<br>.49<br>.93 | Block 2 vs Block 1- p=.83 (two-tailed)<br>Block 3 vs Block 1 - p=.032 (two-tailed)<br>Block 4 vs Block 1 - p <.001 (two-tailed) |
| 1E-Middle | Mixed effects | **Block: F(3,21)=12.04**<br>Session: F(1,7)=1.20<br>Interaction: F(3,17)=.10 | **<.001**<br>.31<br>.95 | Block 2 vs Block 1- p=.99 (two-tailed)<br>Block 3 vs Block 1 - p=.039 (two-tailed)<br>Block 4 vs Block 1 - p <.001 (two-tailed) |
| 1E-Bottom | Mixed effects | Block: F(3,21)=.85<br>Session: F(1,7)=.24<br>Interaction: F(3,15)=1.81 | .48<br>.64<br>.19 |  |
| 1F-Top | RM ANOVA | Block: F(3,21)=1.07<br>Session: F(1,7)=1.31<br>Interaction: F(3,21)=1.05 | .38<br>.29<br>.39 |  |
| 1F-Middle | Mixed effects | Block: F(3,21)=1.06<br>Session: F(1,7)=1.81<br>Interaction: F(3,20)=.93 | .38<br>.22<br>.44 |  |
| 1F-Bottom | RM ANOVA | Block: F(3,21)=.30<br>**Session: F(1,7)=6.26**<br>Interaction: F(3,21)=.568 | .82<br>**.04**<br>.64 | Extinction 1 > Extinction 2 |
| 1G-top | RM ANOVA | **Session: F(4,28)=9.38** | **<.001** | Safe vs Session 1- p=.006 (two-tailed)<br>Safe vs learned - p=.0003 (two-tailed)<br>Safe vs extinction 1 or 2 - p>.94 (two-tailed) |
| 1G-middle | RM ANOVA | **Session: F(4,28)=11.84** | **<.001** | Safe vs Session 1- p=.01 (two-tailed)<br>Safe vs learned - p<.001 (two-tailed)<br>Safe vs extinction 1 or 2 - p>.96 (two-tailed)<br>Learned vs extinction 1 or 2 - p <.01 (two-tailed) |
| 1G-bottom | RM ANOVA | Session: F(4,28)=0.39 | .80 |  |
| Supplement 1A | RM ANOVA | Session: F(2,14)=.72 | .50 |  |
| Supplement 1B | RM ANOVA | Session: F(2,14)=.62 | .55 |  |
| Supplement 1C | RM ANOVA | Session: F(2,14)=1.38 | .28 |  |
| 2A | RM ANOVA | Block: F(3,21)=.63 | .598 |  |
| 2B | RM ANOVA | **Block: F(3,21)=5.69** | **.005** | Block 1 vs 2 - p=.044 (two-tailed)<br>Block 1 vs 3 - p=.002 (two-tailed)<br>Block1 vs 4 - p=.01(two-tailed) |
| 2C | RM ANOVA | Block: F(3,21)=.91 | .45 |  |
| 2D | RM ANOVA | Block: $F(3,21)=1.96$ | .15 | |
| 2E | RM ANOVA | Block: $F(3,21)=.402$ | .75 | |
| 2F | RM ANOVA | Block: $F(3,21)=.214$ | .88 | |
| Supplement 2 (left) | One-sample t test or Wilcoxon signed-rank (IOFC) | Seek: $W=-20$<br>Take: $t(7)=1.92$<br>Reward: $t(7)=1.026$ | .19<br>.095<br>.34 | |
| Supplement 2 (right) | One-sample t test (PL) | **Seek: $t(7)=4.5$**<br>Take: $t(7)=.80$<br>**Reward: $t(7)=2.49$** | **.003**<br>.44<br>**.042** | |
| 3A | Mixed effects | Block: $F(3,20)=.12$ | .94 | |
| 3B | Mixed effects | Block: $F(3,18)=1.19$ | .34 | |
| 3C | Mixed effects | Block: $F(3,27)=.26$ | .85 | |
| 3D | Mixed effects | **Block: $F(3,18)=3.199$** | **.048** | Block 1 vs 2 – $p=.41$ (two-tailed)<br>Block 1 vs 3 – $p=.04$ (two-tailed)<br>Block 1 vs 4 – $p=.057$ (two-tailed) |
| 3E | Mixed effects | Block: $F(3,20)=.12$ | .95 | |
| 3F | Mixed effects | Block: $F(3,18)=.89$ | .46 | |
| 3G | Mixed effects | **Block: $F(3,20)=4.78$** | **.01** | Block 1 vs 2 – $p=.39$ (two-tailed)<br>Block 1 vs 3 – $p=.45$ (two-tailed)<br>Block1 vs 4 – $p=.004$ (two-tailed) |
| 3H | Mixed effects | **Block: $F(3,18)=3.71$** | **.031** | Block 1 vs 2 – $p=.85$ (two-tailed)<br>Block 1 vs 3 – $p=.38$ (two-tailed)<br>Block1 vs 4 – $p=.013$ (two-tailed) |
| 3I | Mixed effects | Block: $F(3,20)=1.276$ | .31 | |
| 3J | Mixed effects | Block: $F(3,18)=.69$ | .57 | |
| 3K | Mixed effects | Block: $F(3,20) = 1.84$ | .17 | |
| 3L | Mixed effects | Block: $F(3,18) = 1.59$ | .23 | |
| 4A | Mixed effects | Shock #: $F(5,23)=1.3$ | .297 | |
| 4B | Mixed effects | Shock #: $F(5,20)=.41$ | .83 | |
| 4C | RM ANOVA | Shock #: $F(5,35)=1.05$ | .40 | |
| 4D | Mixed effects | Shock #: $F(5,23)=.85$ | .53 | |
| 4E | Mixed effects | Shock #: $F(5,20)=.24$ | .938 | |
| 4F | RM ANOVA | Shock #: $F(5,35)=1.07$ | .39 | |
| 5A | RM ANOVA | Block: $F(3,21)=2.46$ | .09 | |
| 5B | RM ANOVA | Block: $F(3,21)=1.63$ | .21 | |
| 5C | RM ANOVA | Block: $F(3,21)=1.58$ | .22 | |
| 5D | RM ANOVA | Block: $F(3,21)=2.69$ | .07 | |
| 5E | RM ANOVA | **Block: $F(3,21)=6.91$** | **.002** | Block 1 vs 2 $p = .009$<br>Block 1 vs 3 $p = .042$<br>Block 1 vs 4 $p = .53$ |
| 5F | RM ANOVA | Block: $F(3,21)=.36$ | .78 | |
| 5G | RM ANOVA | Block: $F(3,21)=1.04$ | .39 | |
| 5H | RM ANOVA | Block: $F(3,21)=0.8$ | .51 | |
| 5I | RM ANOVA | Block: $F(3,21)=.96$ | .43 | |
| 5J | RM ANOVA | Block: $F(3,21)=1.11$ | .37 | |
| 5K | RM ANOVA | Block: $F(3,21)=0.59$ | .63 | |
| 5L | RM ANOVA | Block: $F(3,21)=1.48$ | .25 | |
| 6A-B - Seek | Two way<br>RM AOV | Session: $F(4,28)=.65$<br>Region: $F(1,7)=1.6$<br>**Interaction: $F(4,28)=3.36$** | .63<br>.23<br>**.023** | Comparing each Session to the Safe session for each region<br>PL: session 1 and learned $p < .015$ (one tailed). Extinction 1 and 2 $> .78$.<br>IOFC: all sessions $> .65$ |
| 6A-B - Take | Two way<br>RM AOV | Session: $F(4,28)=1.16$<br>Region: $F(1,7)=.71$<br>**Interaction: $F(4,28)=4.87$** | .35<br>.43<br>**.004** | Comparing each Session to the Safe session for each region<br>PL: all sessions $> .32$<br>IOFC: learned = $.007$ (one tailed). All other sessions $> .32$ |
| 6A-B - Reward | Two way<br>RM AOV | Session: $F(4,28)=.37$<br>Region: $F(1,7)=.50$<br>Interaction: $F(4,28)=1.16$ | .82<br>.50<br>.34 | |
| 6C - Shock | Two way<br>RM AOV | Session: $F(2,14)=3.18$<br>**Region: $F(1,7)=6.34$**<br>Interaction: $F(2,14)=1.16$ | .07<br>**.039**<br>.34 | PL>IOFC |
| 7F-left | Two way<br>AOV | **Block: $F(3, 2644)=8.59$**<br>**Session: $F(4, 2644)=12.03$**<br>**Interaction: $F(12, 2644)=6.54$** | **<.001**<br>**<.001**<br>**<.001** | Comparing each block to the block in the Safe session<br>Block 1: all $p$ values $> .14$<br>Block 2: learned vs safe = $.058$, all other $p$ values $> .18$<br>Block 3: learned vs safe = $.001$, extinction 1 vs safe = $.066$; all other $p$ values $> .51$<br>Block 4: session 1 vs safe = $.001$; learned vs safe = $.001$, extinction 1 vs safe = $.036$; extinction 2 = $.246$ |
| 7F-right | Two way<br>AOV | Block: $F(3,2644)=.269$<br>Session: $F(4,2644)=1.96$<br>Interaction: $F(12,2644)=1.237$ | .83<br>.10<br>.25 | |

**Supp. Figure 1:**
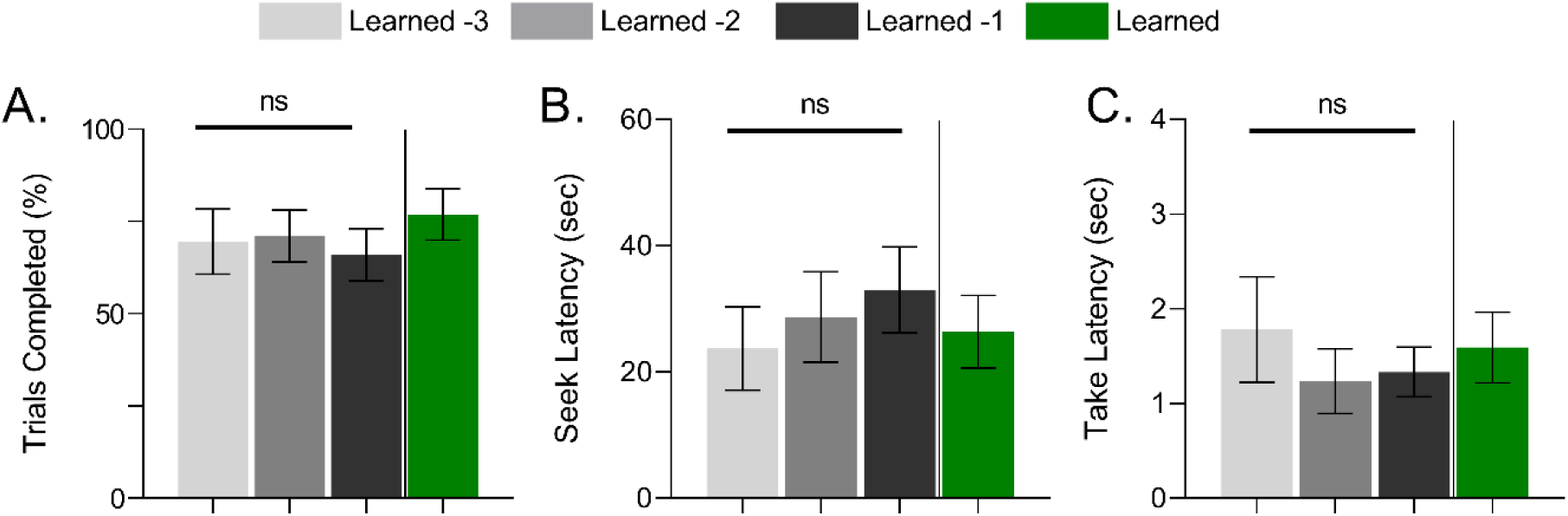
Percent of trials completion or action latency over risk blocks in the Learned session (green – Learned) or the 3 sessions preceding recording (shades of grey – Learned-X). **A.** Mean ± SEM trial completion over sessions. **B.** Mean ± SEM seek action latency over sessions. **C.** Mean ± SEM take action latency over sessions. n=8 rats, ns= no significant difference over Learned –X sessions.

**Supp. Figure 2:**
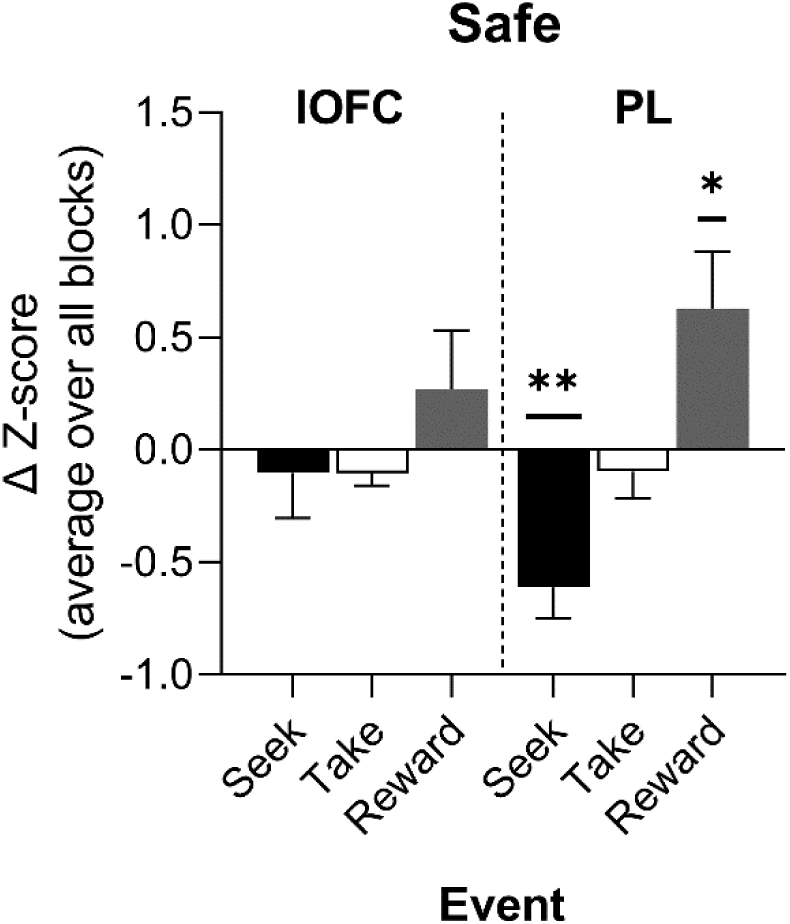
Block averaged change scores of neuronal calcium activity in lOFC or PL during seek, take, or reward events over the Phase I Safe session. Mean lOFC (left bars) and mean PL (right-bars) response change scores in Safe sessions of Phase I. n=8 rats. * p *<* .05 or ** p *<* .01 versus 0, one sample t-test. Data are presented as mean ± SEM.

